# *broad* controls leg imaginal disc morphogenesis in Drosophila via regulation of cell shape changes and remodeling of extracellular matrix

**DOI:** 10.1101/2019.12.21.885848

**Authors:** Clinton Rice, Stuart Macdonald, Xiaochen Wang, Robert E Ward

## Abstract

Imaginal disc morphogenesis during metamorphosis in *Drosophila melanogaster* provides an excellent model to uncover molecular mechanisms by which hormonal signals effect physical changes during development. The *broad* (*br*) *Z2* isoform encodes a transcription factor required for disc morphogenesis in response to 20-hydroxyecdysone, yet how it accomplishes this remains largely unknown. Here, we show that amorphic *br^5^* mutant discs fail to remodel their basal extracellular matrix (ECM) after puparium formation and do not undergo necessary cell shape changes. RNA sequencing of wild type and mutant leg discs identified 717 genes differentially regulated by *br*; functional studies reveal that several are required for adult leg formation, particularly those involved in remodeling the ECM. Additionally, *br Z2* expression is abruptly shut down at the onset of metamorphosis, and expressing it beyond this time results in failure of leg development during the late prepupal and pupal stages. Taken together, our results suggest that *br Z2* is required to drive ECM remodeling, change cell shape, and maintain metabolic activity through the mid prepupal stage, but must be switched off to allow expression of pupation genes.

**Summary Statement:** The *Drosophila melanogaster* ecdysone-responding transcription factor *broad* controls morphogenetic processes in leg imaginal discs during metamorphosis through regulation of genes involved in extracellular matrix remodeling, metabolism, and cell shape changes and rearrangements.

## Introduction

Tissue morphogenesis is required for the elaboration of the body axis and organs during metazoan development. In some contexts, hormonal signals provide temporal cues to coordinate these morphogenetic events. Imaginal disc morphogenesis in the fruit fly *Drosophila melanogaster* provides an excellent model to uncover molecular mechanisms by which hormonal signals are translated into the physical changes that occur during development. Imaginal discs are diploid tissues found within the larva that give rise to the adult integument during metamorphosis (Willis, 1974; Csaba, 1977; Fristrom, 1988). Both classical experiments and more recent live imaging studies (De las Heras *et al*., 2018; Diaz-de-la-Loza *et al*., 2018) have revealed requirements for cell shape changes and rearrangements, as well as for remodeling of the extracellular matrix (ECM) in imaginal disc morphogenesis during metamorphosis. These experiments also demonstrate the key role that 20-hydroxyecdysone (hereafter referred to as ecdysone) plays in directing these processes. Ecdysone acts through a transcriptional cascade— the hormone binds to its heterodimeric receptor, which acts as a DNA-binding activator to drive transcription of early-response genes, including *Eip74EF*, *Eip75B*, and *broad* (*br*), which themselves encode DNA-binding proteins that activate late-response genes (Chao and Guild, 1986; Feigl *et al*., 1989; Janknecht *et al*., 1989; Burtis *et al*., 1990; Segraves and Hogness, 1990; DiBello *et al*., 1991; Yao *et al*., 1993; Crossgrove *et al*., 1996). Despite extensive research into *Drosophila* metamorphosis, parts of the pathway connecting the ecdysone cue to the effectors involved in imaginal disc morphogenesis have yet to be elucidated.

Of the ecdysone signaling early-response genes, *br* appears to have the most direct effects on imaginal disc morphogenesis (Kiss *et al*., 1988). *br* encodes four transcription factors, each carrying a unique zinc finger domain spliced to a common Broad-Complex, Tramtrack, and Bric a Brac/Pox virus and Zinc finger (BTB/POZ) domain (DiBello *et al*., 1991; von Kalm *et al*., 1994; Bayer *et al*., 1996). These four isoforms, known as the Z1, Z2, Z3, and Z4 isoforms, have three genetically separate functions (*br*, *reduced bristles on the palpus* and *2Bc*), with the Z2 isoform performing the classic *br* function (Bayer *et al*., 1997). This function is critical for imaginal disc morphogenesis: animals lacking functional Br Z2 have discs that fail to elongate; these animals die during the prepupal stage prior to head eversion (Kiss *et al*., 1988). The various isoforms of Br also add a layer of complexity to the regulation of metamorphic genes. *br* does not simply respond to the ecdysone cue through a general increase in transcription that subsequently effects the increased transcription of late-response genes; rather, the isoforms exhibit a dynamic pattern of expression that differs by tissue (Huet *et al*., 1993). In the imaginal discs, the expression of the Z2 transcript rises dramatically approximately 4 hours before pupariation, but greatly decreases in the hours after pupariation, while the expression of the Z1 transcript greatly increases between 2 and 4 hours after pupariation (Emery *et al*., 1994; Bayer *et al*., 1996). This late larval pulse of Z2 expression is critical for activation of late-response genes; however, the identities of these genes remain largely unknown. The illumination of the link between the ecdysone cue and morphogenetic effects in imaginal discs hinges upon the identification of these late-response genes and how they are regulated by ecdysone and *br*.

Previous screens using the hypomorphic *br^1^* allele identified a number of *br*-interacting genes including *Stubble* (*Sb*), *zipper*, *Rho1*, *Tropomyosin 1*, *blistered*, and *ImpE3* (Beaton *et al*., 1988; Ward *et al*., 2003), although the majority of these genes are not transcriptional targets of *br* (Ward *et al*., 2003). Nevertheless, the identity of these genes suggests roles for *br* in the major processes implicated in hormonal control of imaginal disc morphogenesis, namely cell shape changes/rearrangements and modification of the ECM. Here we show that cell shape changes and cell rearrangements fail to occur normally in *br^5^* mutant prepupae. In addition, consistent with previous work demonstrating that the ECM provides a constraining force and must be degraded to allow disc elongation (Pastor-Pareja and Xu, 2011; Diaz-de-la-Loza *et al*., 2018; Maria-del-Carmen *et al*., 2018), we show that the basal ECM protein Collagen IV is not substantially degraded in leg imaginal discs from amorphic *br^5^* mutant animals as old as 8 hours after puparium formation (APF). In this study, we also use this allele to identify specific genes regulated by *br* in the leg discs at the onset of metamorphosis through an RNA sequencing-based approach. This approach identified over 700 *br*-regulated genes, including genes with known metabolic and developmental functions, including ECM organization and modification. We tested a subset of these genes for roles in leg morphogenesis through RNAi and found several that are necessary for proper development of the adult legs. These results demonstrate the value of a transcriptional target-based approach to identifying morphogenetic genes, and suggest that *br* regulates morphogenesis through the regulation of genes involved in multiple critical processes.

## Results

### Phenotypic characterization of *br^5^* mutant leg imaginal discs

In order to identify genes regulated by *br* during metamorphosis using an RNA sequencing (RNA-Seq) approach, we wanted to use a null *br* allele. The *br^5^* allele has been reported to be amorphic, with homozygous mutants failing to develop past the early prepupal stage (Kiss *et al*., 1988). Consistent with this notion, *br^5^* encodes a protein with a His^492^–Tyr substitution in the conserved Z2 zinc finger domain, suggesting that it has defective DNA binding properties (personal communication from Laurie von Kalm and confirmed in the RNA sequencing; see materials and methods). We previously used live imaging to show that *br^5^* hemizygous mutants failed to complete metamorphosis (Ward *et al*., 2003). To more closely examine how imaginal disc development is disrupted in these animals, we dissected and compared leg discs from *br^5^* and *w^1118^* larvae and prepupae (Fig. 1). *br^5^* leg discs resemble *w^1118^* leg discs during larval stages (through 4 hours before pupariation), consisting of a columnar epithelium with folds that make 3-4 concentric circles. The discs bulge from the central circles (which develop into the distal-most leg segments), and are covered by the peripodial epithelium. By the prepupal stage, leg discs begin to differ between the two genotypes. Both *br^5^* and *w^1118^* discs begin to elongate in a telescoping fashion at the onset of metamorphosis (0 hours), beginning with the centermost regions. However, by two hours after pupariation, elongation of *br^5^* discs clearly lags behind that of *w^1118^* discs. *w^1118^* discs continue to elongate and by 4 hours after pupariation, the future 5 tarsal segments and distal tibia are clearly identifiable underneath the peripodial epithelium. *br^5^* mutant discs, on the other hand, form shorter, wider structures with deeper folds between the tarsal segments. *br^5^* mutant discs show only limited elongation over subsequent hours and fail to evert to the outside of the prepupa. Notably, imaginal discs can be found inside degenerating *br^5^* mutant late prepupae that look similar to +4 hour mutant discs, including having an intact peripodial epithelium (Fig. 1K).

**Fig. 1.**
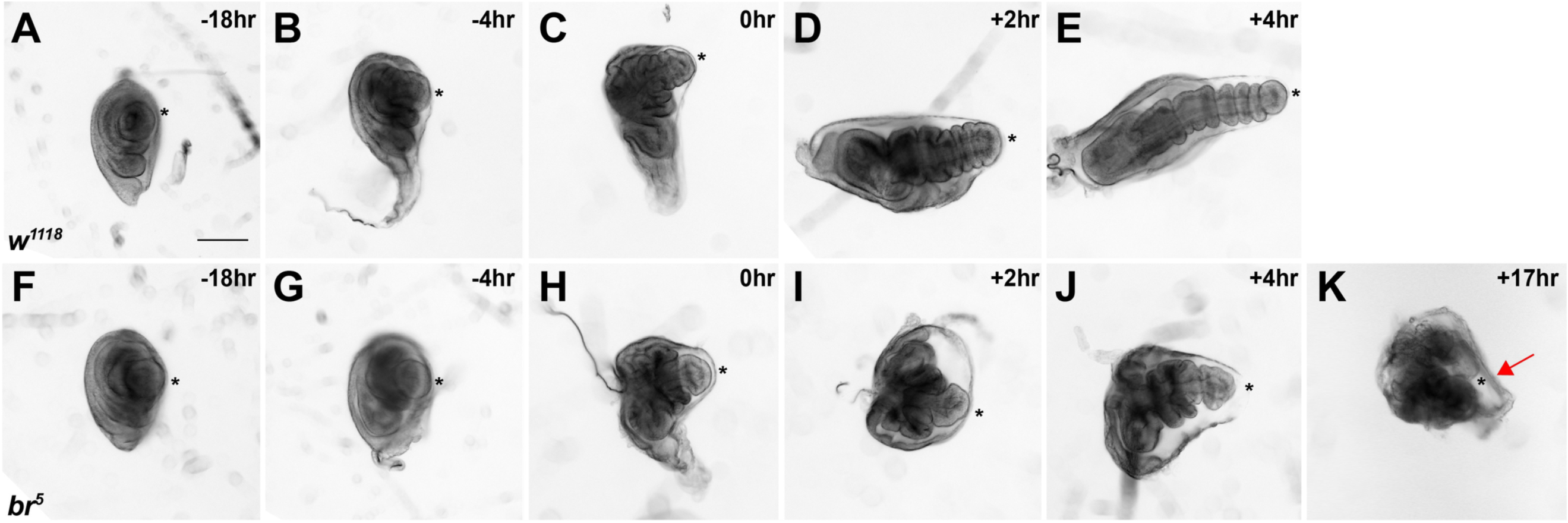
*Broad* is required for normal prepupal leg imaginal disc development. Brightfield photomicrographs of leg imaginal discs from *w^1118^* (A-E) and *br^5^* (F-K) mutant larvae and prepupae. Times given are relative to puparium formation. Note that leg imaginal discs are similar between *w^1118^*and *br^5^* during larval time points (−18 hr and −4 hr), but are noticeably different starting at 0 hr. Whereas *w^1118^* discs elongate from the center (distal tarsal segment; indicated by asterisk) and are segmented by +4 hr, *br^5^* discs show limited elongation and have wider tarsal segments. Intact leg imaginal discs can be found inside dead *br^5^* prepupae as late as 17 hr after pupariation. These late imaginal discs have not elongated much past the +4 stage and often have an intact peripodial epithelium (arrow). Scale bar equals 100 um.

The disruption of prepupal elongation in *br^5^* mutants, coupled with the presence of the peripodial epithelium in late-staged prepupae (which is known to be degraded by matrix metalloproteinases (Proag *et al*., 2019)) motivated us to examine ECM breakdown in wild type and *br^5^* mutant discs. We therefore crossed a GFP-tagged version of Collagen IV (Vkg-GFP) into *w^1118^* and *br^5^* and examined their leg discs at various time points during metamorphosis (Fig. 2). At early stages (0 and 2 hours after pupariation), wild type and *br^5^* leg discs closely resemble one another, with fibrous cylindrical or nearly-conical basal ECM structures. By 4 hours after pupariation, some wild type discs exhibit degradation of the basal ECM. This degradation takes the form of a “clearing” of a central channel along the length of the disc. In total, 37.5% of wild type discs (*n* = 32, from 12 animals) showed this clearing at 4 hours after pupariation, 66.7% (*n* = 21, from 12 animals) at 6 hours, and 100% (*n* = 12, from 8 animals) at 7 hours. In contrast, most *br^5^* mutant discs retained the fibrous ECM appearance as late as 8 hours after pupariation (Fig. 2J). Only 12.0% of discs (*n* = 25, from 7 animals) showed partial clearing; the other discs showed no clearing.

**Fig. 2.**
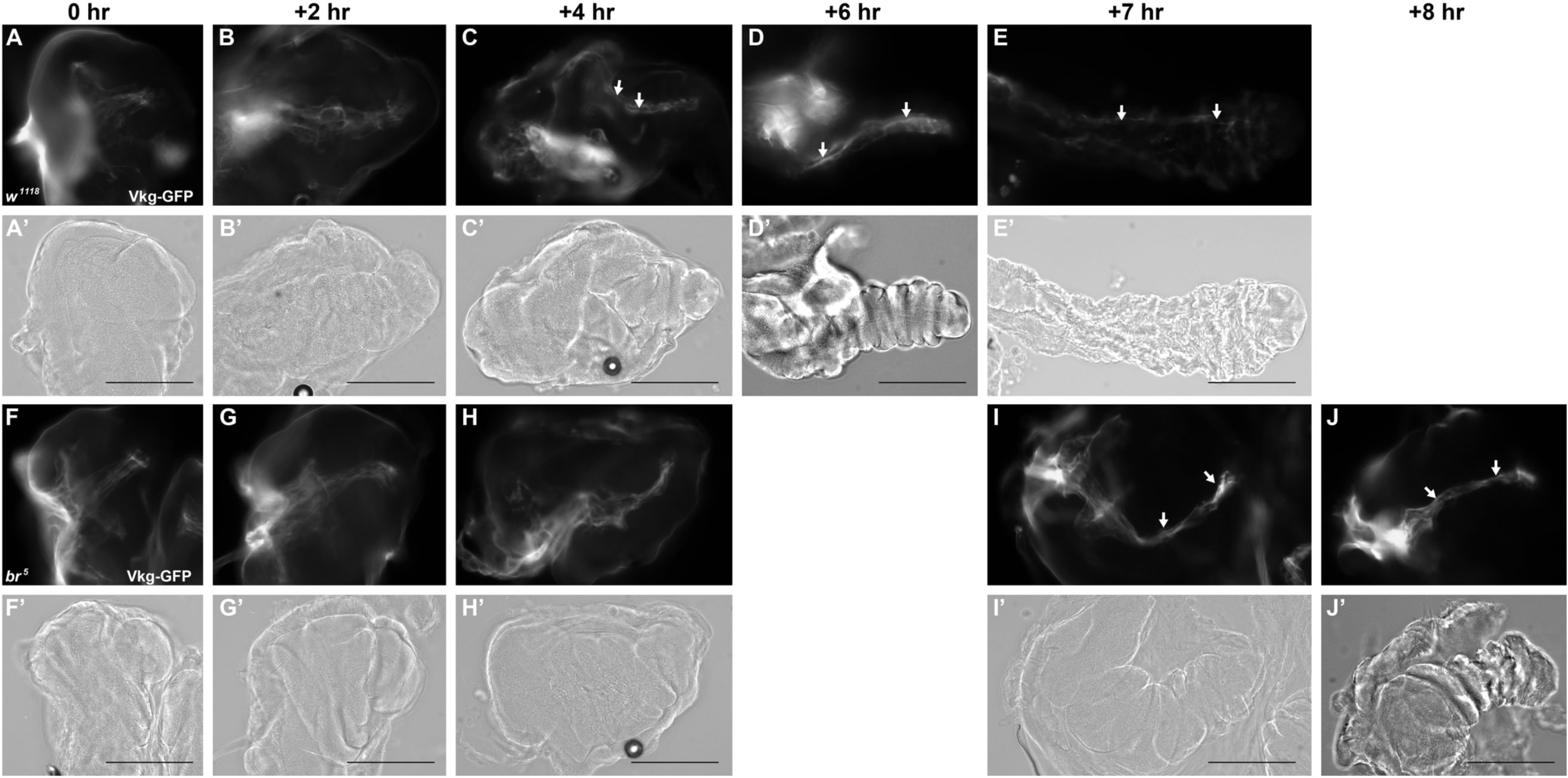
*broad* is required for efficient degradation of the basal ECM in prepupal leg imaginal discs. Wide-field fluorescence (A-J) and Brightfield (A’-J’) photomicrographs (at same focal plane) of leg imaginal discs from *w^1118^* (A-E) and *br^5^* mutant (F-J) prepupae expressing Viking-GFP (Collagen IV). Ages of the animals relative to puparium formation are indicated above the figures. Note that Vkg-GFP forms cables of ECM lining the lumen at the basal side of elongating leg discs (arrows), which are gradually cleared from +4 to +7 hours APF in *w^1118^* leg discs. Similar clearing is not observed in the *br^5^* mutant discs and persists at least through 8 hours APF (J) in many animals. Scale bar equals 100 um.

Since *br^5^* mutant leg imaginal discs show reduced elongation and defective proteolysis of the ECM, we wondered whether an exogenous protease could restore normal elongation to the *br^5^* discs. We therefore dissected 3 leg imaginal discs from individual *w^1118^* or *br^5^* animals at the onset of metamorphosis and treated one with 0% trypsin (PBS control), another with 0.0025% trypsin and the third with 0.025% trypsin for 15 minutes at room temperature. Leg imaginal discs from 18 of the 21 *w^1118^* animals had clearly elongated after 15 minutes in 0.0025% trypsin (Fig. 3B). The tarsal segments in these discs were still obviously segmented, although in some cases the depth of the folds at these segment boundaries was reduced relative to the PBS control discs. *w^1118^* leg discs incubated for 15 minutes in 0.025% trypsin showed even greater elongation (in 100% of the discs), but only with noticeable tarsal segment boundaries in 2 of the discs, whereas the remainder had elongated discs with the tarsal segments appearing as one long continuous segment (Fig. 3C). In the *br^5^* mutant leg discs, there was a clear difference in the morphology of the leg discs at the onset of the experiment (which remained unchanged through 15 minutes in PBS, data not shown) compared to the *w^1118^* discs, in which the distal tarsal segment was rounder and the folds between the tarsal segments were noticeably deeper (Fig. 3D vs 3A). 10 of the 17 discs treated with 0.0025% trypsin showed some elongation of the tarsal segments, but no shallowing of the segment boundaries (Fig. 3E). Similarly, 12 of the 17 discs treated with 0.025% trypsin showed an increase in the degree of elongation. The morphology of these discs was clearly distinct from *w^1118^*treated similarly. The absolute level of elongation in *br^5^* was less than in *w^1118^*, and more interestingly the *br^5^* discs did not eliminate the folds between segments, but rather deepened those folds (Fig. 3F). This morphology was observed in 14 of the 17 discs examined.

**Fig. 3.**
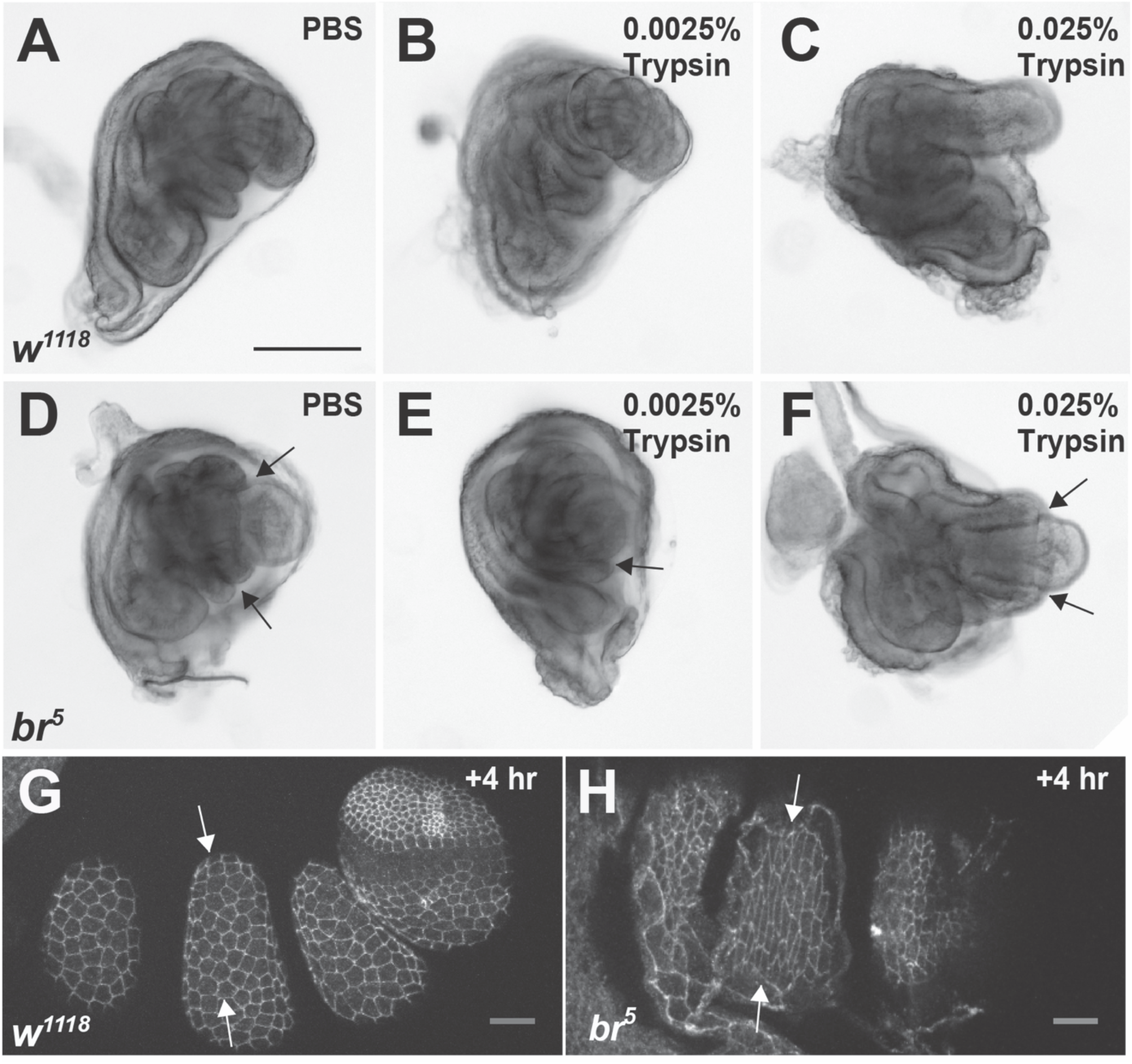
*br^5^* leg discs show aberrant elongation upon application of trypsin, and retain anisometric cell shapes through 4 hours after pupariation. (A-F) Brightfield photomicrographs of leg imaginal discs from a representative *w^1118^*(A-C) and *br^5^* (D-F) mutant 0 hr prepupa incubated for 15 minutes in PBS (A and D) or 0.0025% trypsin (B and E) or 0.025% trypsin (C and F). All three discs are from the same animal. The *w^1118^* discs elongate in the lower dose of trypsin, and elongate more in the higher dose of trypsin to the point where the segmental folds are pulled smooth. In contrast, *br^5^* mutant discs show less elongation in trypsin overall, and deeper folds in the presumptive tarsal segments (arrows). (G, H) Confocal optical sections of tarsal segments from leg imaginal discs from *w^1118^* (G) and *br^5^* (H) mutant +4 hr leg imaginal discs stained with antibodies against DE-Cadherin (distal tarsal segments are to the right in each image). Note that in the *w^1118^* discs, epithelial cells in the tarsal segments are mostly isometric, whereas cells in the *br^5^* tarsal segments are anisometric with longer circumferential cell lengths (arrows). Scale bar equals 100 um.

Given that trypsin was not able to elongate *br^5^* leg discs to the extent of that observed in *w^1118^* discs, and that *br^5^* discs display aberrant morphology as early as the onset of metamorphosis, we suspected that the underlying cell shape changes and/or rearrangements that normally occur in wild type animals do not occur in these mutant animals. To address this question, we examined cell shapes in the distal tarsal segments of +4 hr prepupal leg imaginal discs from *w^1118^* and *br^5^* mutant animals using antibodies against DE-Cadherin. Whereas epidermal cells from *w^1118^* +4hr leg discs are generally isometric, being equally long in the proximal-distal and circumferential dimensions (Fig. 3G), epidermal cells from *br^5^* +4hr leg discs show considerable anisometry, with longer cell dimensions in the circumferential axis than the proximal-distal axis (Fig. 3H). This phenotype is completely penetrant (based upon 5 experiments with leg discs from more than 30 mutant animals). The *br^5^* mutant distal tarsal segments are also wider at +4 hours than *w^1118^* leg discs (compare Fig. 1E vs 1J), suggesting a defect in cell rearrangements that are known to occur in wild type (Condic *et al*., 1991), although the defect in segmentation in the *br^5^* discs make this difficult to quantify.

### Identification and bioinformatic characterization of genes regulated by *br* in prepupal leg discs

In order to better understand the role of *broad* in regulating imaginal leg morphogenesis, we used RNAseq to identify genes that are differentially expressed between *w^1118^* and *br^5^*. We dissected leg discs from these animals at the onset of metamorphosis (0 hr), the time we first see differences in the tissues between genotypes. We generated libraries from three biological replicates per genotype and sequenced them on an Illumina HiSeq 2500. We then mapped reads to the reference genome (release 5.3) with TopHat, and used the Cufflinks pipeline to identify a total of 717 genes that were significantly differentially expressed between the two genotypes (at a False Discovery Rate (FDR) of 5%) (Fig. 4) (Trapnell *et al*., 2009; Trapnell *et al*., 2010; Roberts *et al*., 2011; Kim *et al*., 2013; Trapnell *et al*., 2013). About two-thirds of these genes (66.9%) were more highly expressed in the *br^5^* genotype. We refer to these genes as *br*-repressed genes (and genes that were more expressed in *w^1118^* as *br*-induced genes). We used *br*-induced and *br*-repressed genes with fold changes ≥ 1.5 and FPKM ≥ 5 in at least one of the samples (leaving 155 *br*-induced genes and 356 *br*-repressed genes) to identify significant gene ontology (GO) terms using the Gene Ontology Consortium’s GO Enrichment Analysis (The Gene Ontology Consortium. 2015) (Table S1). Many *br-*induced genes have predicted metabolic functions: significantly enriched GO terms for this set of genes included “glycolytic process,” “glycogen metabolic process,” and “carbohydrate biosynthetic process” (Table 1 and Table S2). KEGG (Kyoto Encyclopedia of Genes and Genomes) pathway analysis of the same set of genes using WebGestalt (Wang *et al*., 2017) revealed significant overrepresentation of genes involved in Glycolysis/Gluconeogenesis, and several sugar metabolism pathways for *br-*induced genes, and genes involved in several amino acid metabolism pathways in *br-*repressed genes (Table S3). In addition, several GO terms that were significantly overrepresented in *br*-induced genes are related to cell signaling and development including “Notch signaling,” “Molting cycle, chitin-based cuticle,’ and “Developmental process” (Table 1).

**Fig. 4.**
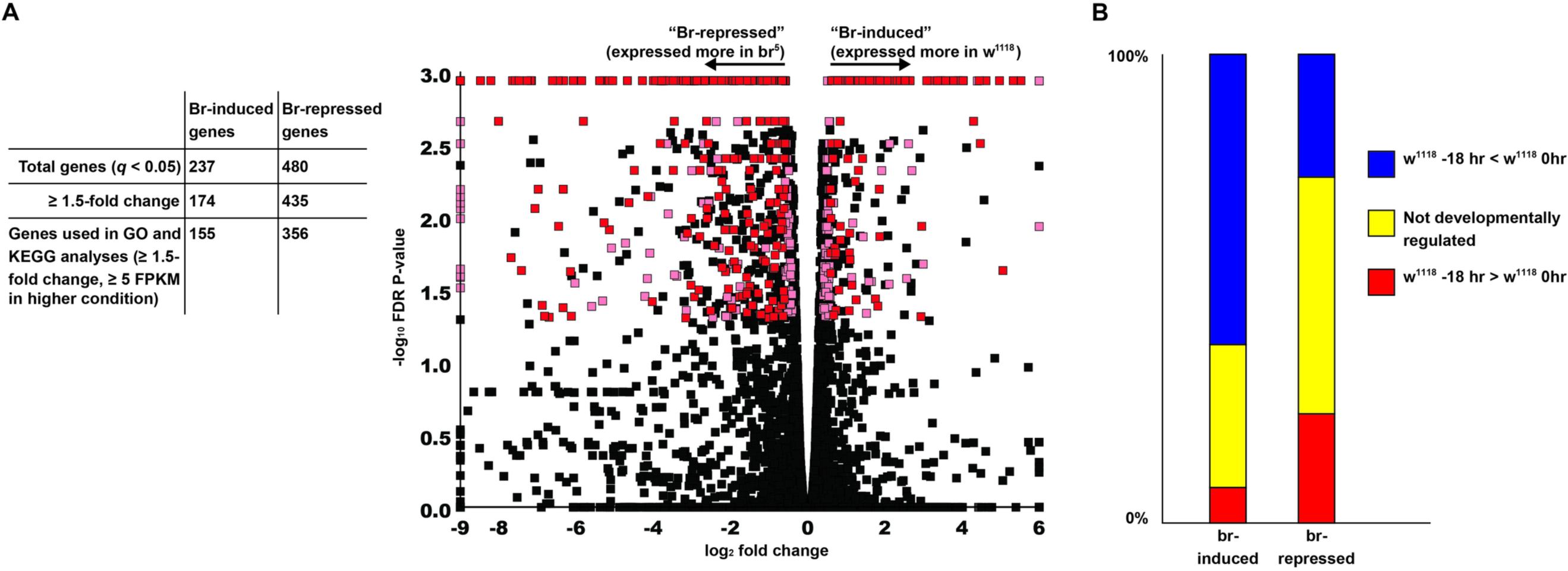
Over 700 genes are significantly differentially expressed in the absence of functional *br*-*Z2*. (A) Volcano plot of genes differentially regulated in *br^5^* mutant leg imaginal discs relative to *w^1118^* controls. Red denotes genes that were used in the GO and KEGG analyses, while pink denotes genes that were significantly differentially regulated but not included in the analyses (fold change < 1.5 or FPKM < 5). (B) Overlap of *br*-regulated genes with developmentally regulated genes. Most *br*-induced genes were developmentally upregulated (higher in 0 hr *w^1118^* prepupae than in −18 hr *w^1118^* larvae), while relatively few *br*-repressed genes were developmentally downregulated.

**Table 1:** Selected significantly enriched biological process GO terms for br-induced genes.

| GO Term | # of genes | Expected | Fold change | Significance |
| --- | --- | --- | --- | --- |
| single-organism carbohydrate catabolic process | 9 | 0.27 | 32.93 | 3.91E-08 |
| ATP generation from ADP | 8 | 0.23 | 35.37 | 3.14E-07 |
| glycolytic process | 8 | 0.23 | 35.37 | 3.14E-07 |
| ADP metabolic process | 8 | 0.25 | 32.65 | 5.86E-07 |
| pyruvate metabolic process | 8 | 0.33 | 24.25 | 5.88E-06 |
| Notch signaling pathway | 8 | 0.52 | 15.43 | 1.87E-04 |
| molting cycle, chitin-based cuticle | 8 | 0.91 | 8.75 | 1.26E-02 |
| developmental process | 55 | 31.49 | 1.75 | 1.39E-02 |
| anatomical structure development | 53 | 30.32 | 1.75 | 2.30E-02 |
| organ morphogenesis | 22 | 7.98 | 2.76 | 4.11E-02 |

Since *br* serves as an early-response gene to the late larval ecdysone pulse (Chao and Guild, 1986; DiBello *et al*., 1991), we expected that *br*-regulated genes would essentially represent a subset of genes regulated by ecdysone. To test this idea, we conducted a second set of RNA-seq analyses comparing genes expressed in mid third instar (−18 hr) *w^1118^* leg imaginal discs to those expressed in *w^1118^*leg imaginal discs at the onset of metamorphosis (0 hr). A total of 1,186 genes are significantly upregulated at the 0 hr stage relative to the −18 hr stage (developmentally-induced genes), while 1,193 genes are significantly downregulated (developmentally-repressed genes) (Table S4). We anticipated that these developmentally-regulated genes would serve as a proxy for ecdysone-regulated genes; consistent with this expectation, many known ecdysone-regulated genes are found in this data set including *Eip74EF*, *E23*, and *Cyp18a1* (Janknecht *et al*., 1989; Burtis *et al*., 1990; Hurban and Thummel, 1993; Bassett *et al*., 1997; Hock *et al*., 2000). Significantly enriched GO terms in the developmentally-induced genes include numerous terms related to developmental processes, such as “imaginal disc eversion” and “regulation of tube size”, while terms relating to mitosis and DNA replication are significantly overrepresented in the developmentally-repressed genes (Table S5). Of the 237 genes induced by *br,* 147 (62.0%) were upregulated at the 0 hr stage in *w^1118^* flies (Fig. 4B). In contrast with the predictions of the hierarchical model of ecdysone-driven gene regulation, more *br*-repressed genes were upregulated in 0 hr prepupae than downregulated, although the difference was not significant (126 genes and 112 genes, respectively) (Fig. 4B).

The absence of metabolic GO terms among those enriched in the developmentally-induced genes suggests that wild type discs persist in their metabolic activity even as the larval tissues are down-regulating metabolism (Merkey, 2011), and that this maintenance of metabolism is dependent upon *br* function. This notion is consistent with the observation that *br* mutant leg discs appear to arrest development at about 4 hours APF (Fig. 1). To address this experimentally, we quantified ATP levels in +4 hr *w^1118^* and *br^5^* mutant mixed leg and wing imaginal discs using a luciferase-based assay. To our surprise, we found that ATP levels were significantly higher in *br^5^* discs (P=0.03, Fig. S1).

### Change in *br* expression in imaginal discs features an isoform switch at metamorphosis

Although the large proportion of *br*-regulated genes that are temporally regulated at metamorphosis is generally consistent with the hierarchical model of *br* gene regulation, in which *br* is induced by ecdysone and in turn induces the late response genes, we also found many *br*-regulated genes that do not appear to be developmentally-regulated (Fig. 4B). We therefore suspected a more complex relationship between ecdysone-regulated and *br*-regulated genes. Given that previous work has shown differential expression of *br* isoforms in different tissues (Huet *et al*., 1993), we wanted to investigate the expression of *br* in imaginal discs relative to the late larval ecdysone pulse. We therefore performed a northern blot analysis using total RNA isolated from whole animals and from combined leg and wing discs at four time points from 18 hours before pupariation to 4 hours after pupariation (Fig. 5A), and probed the samples for expression of all four *br* isoforms (using a probe specific to the common BTB/POZ domain). In both whole animals and imaginal discs, the *br* Z2 and/or Z3 isoforms are visible at −18, −4, and 0 hr, although these appear to be less highly expressed at −18 hr in whole animals, while their expression begins to decrease by 0 hr in the discs. In both whole animals and imaginal discs, expression of the Z2 and/or Z3 isoforms ends before 4 hours after pupariation. In contrast, the Z1 isoform of *br* shows minimal expression 18 hours before pupariation, and highest expression 4 hours after pupariation, particularly in imaginal discs.

**Fig. 5.**
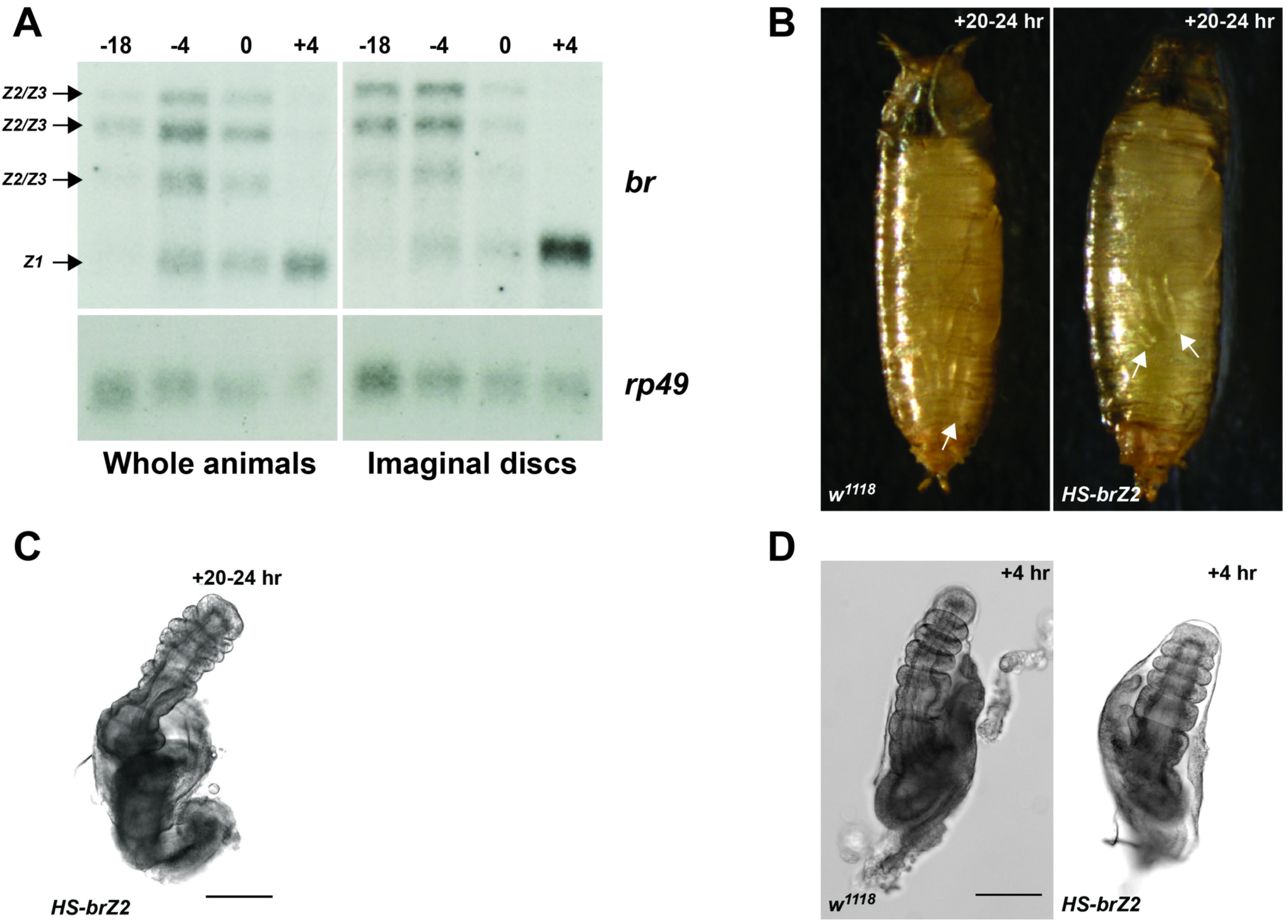
Downregulation of *br* expression at the onset of metamorphosis is required for normal late prepupal and pupal development. (A) Northern blot analysis of total RNA extracted from whole animals or leg and wing imaginal discs around the time of metamorphosis showing expression of *br*. RNA was isolated from larvae at 18 hours and 4 hours before pupariation and from pupae at 0 hours and 4 hours after pupariation. Expression of the *Z2* and/or *Z3* isoforms is substantially reduced by the time of pupariation (0 hr) and these transcripts are absent by 4 hours after pupariation, at which point the *Z1* isoform is highly expressed. *rp49* was used as a control for loading and RNA transfer. (B) Photomicrographs of *w^1118^* (left side) and *HS-brZ2* (right side) early pupae ∼20-24 hr after pupariation that had been heat shocked at 37° for 60 minutes within 6 hours before puparation. The *HS-brZ2* pupae has short legs (arrows indicate the distal tips of legs in the two animals). (C) Brightfield photomicrograph of a leg disc from a *HS-brZ2* dead prepupae ∼20-24 hr after pupariation that had been heat shocked at 37° for 60 minutes within 6 hours before puparation. Note that this animal had not pupated and the leg disc was found inside the degenerating prepupae. This terminal phenotype is similar to what is seen in a wild type prepupae at about 6 or 7 hours after pupariation. (D) Brightfield photomicrograph of leg discs from *w^1118^*(left side) and *HS-brZ2* (right side) 4 hr after pupariation that had been heat shocked at 37° for 60 minutes within 6 hours before puparation. Note that the *HS-brZ2* leg disc looks similar to that from *w^1118^*. Scale bar equals 100 um.

Since transcription of the Z2 isoform of *br* appears to stop at the onset of metamorphosis, we suspected that derepression of some Br-repressed genes may be critical for leg morphogenesis. We therefore attempted to disrupt this process by misexpressing the *br Z2* isoform during the prepupal stage. To accomplish this, we heat shocked late larvae carrying a heat inducible-*br* Z2 construct (Crossgrove *et al*., 1996), collected all animals that pupariated within 6 hours after the heat shock, and followed the animals through prepupal and pupal development. In six separate experiments totaling 99 animals, only 50% of the *HS-brZ2* animals pupated, and of those only 4 developed to late pupae, with the remaining dying as early pupae. In addition, most of those that pupated had short legs and wings (Fig. 5B) or poorly everted heads. In contrast >90% of *w^1118^* animals treated in the same way pupated (n=80), and 73% eclosed. Dissection of leg imaginal discs from the dead *hs-brZ2* prepupae revealed variable terminal leg phenotypes, but most appeared to arrest at stages comparable to wild type legs between 4 and 7 hours APF (Fig. 5C). To confirm that the first several hours of prepupal development occurred normally in these animals, we repeated the experiment, collected animals as they pupariated, aged them four hours and then dissected and imaged the leg imaginal discs. All the legs appeared to have elongated normally at +4 hours regardless of how many hours (0-5 hr) prior to pupariation that they were heat shocked (Fig. 5D). Taken together, these results suggest that either the continued expression of some *br*-induced genes or failure to derepress some *br*-repressed genes are detrimental to later prepupal and pupal leg development.

### Functional analysis of *br*-induced genes in leg morphogenesis

Given the important role that *br* plays in leg disc morphogenesis, we expected that some of the genes induced by *br* in the leg discs at metamorphosis would also be critical for this process. To test this, we induced RNA interference (RNAi) using *Distal-less* (*Dll*)*-GAL4* for 27 *br*-induced genes (Table 2). *Dll-GAL4* is expressed in the distal tibia and tarsal segments during late larval and prepupal time points (Cohen, 1993; Ward *et al*., 2003). We investigated genes with peptidase or peptidase inhibition functions due to our observation that basal ECM degradation is perturbed in *br^5^* mutant leg discs (Fig. 3), and we were particularly interested in *Stubble* (*Sb*) because it has been shown in previous screens to have an enhancer of *br* effect (Beaton *et al*., 1988; Gotwals and Fristrom, 1991; Ward *et al*., 2003). We tested members of the *Enhancer of split* complex due to the enrichment of the “Notch signaling” GO term among *br*-induced genes. We also tested a collection of genes that have chitin-related functions since the GO enrichment showed an overrepresentation of these genes in the *br*-induced set. Finally, we also tested several genes in the Ecdysone-induced 71E cluster (Eig71E), since six of the eleven Eig71E genes were found to be *br*-induced.

**Table 2:** Frequency of leg defects in flies expressing RNAi against br-induced genes.

| Target gene | TRiP or VDRC? | Line # | % carrying *dll>Gal4* | % with malformed legs | Note |
| --- | --- | --- | --- | --- | --- |
| **Protease and protease inhibitors** |  |  |  |  |  |
| *CG3355* | TRiP | 52897 | 48.9% | 0.0% |  |
| *CG5639* | TRiP | 57376 | 48.2% | 0.0% |  |
|  | VDRC | 1306 | 50.0% | 1.2% |  |
| *CG8170* | TRiP | 61889 | <u>36.2%</u> | 0.0% |  |
| ***CG9416*** | **TRiP** | **57563** | **<u>28.6%</u>** | **37.5%** | **Bent tibiae** |
|  | VDRC | 10064 | 45.9% | 2.6% |  |
| ***Stubble*** | **TRiP** | **42647** | **<u>29.8%</u>** | **18.3%** | **Bent femurs, tibiae, and tarsals** |
| *Serine Protease Immune Response Integrator* | TRiP | 42882 | 47.1% | 0.0% |  |
| **Notch pathway genes** |  |  |  |  |  |
| *Enhancer of split m4* | TRiP | 61261 | <u>41.3%</u> | 1.5% |  |
| *Enhancer of split m7* | TRiP | 29327 | 51.6% | 1.3% |  |
| *Enhancer of split m8* | TRiP | 26322 | <u>39.4%</u> | 1.2% |  |
|  | VDRC | 37686 | 45.6% | 0.0% |  |
| *Enhancer of split mα* | VDRC | 35886 | <u>0.0%</u> | N/A | No adult flies—larvae failed to molt |
| *Enhancer of split mγ* | TRiP | 51762 | 48.0% | 0.0% |  |
| *scabrous* | TRiP | 56928 | 54.3% | 0.0% |  |
| **Eig71E genes** |  |  |  |  |  |
| *Ecdysone-induced gene 71Ec* | TRiP | 57384 | 45.9% | 0.0% |  |
| *Ecdysone-induced gene 71Ed* | TRiP | 56952 | 52.4% | 0.0% |  |
| *Ecdysone-induced gene 71Ef* | TRiP | 55930 | 46.1% | 0.0% |  |
| *Ecdysone-induced gene 71Eg* | TRiP | 56953 | 47.9% | 0.9% |  |
| **Chitin related genes** |  |  |  |  |  |
| *dusky* | TRiP | 34382 | 55.3% | 0.0% | All flies had slightly curved tibiae, but not severe enough to be counted as malformed |
|  | **VDRC** | **3299** | **41.5%** | **100.0%** | **Bent tibiae** |
| *Imaginal disc growth factor 4* | TRiP | 55381 | 45.8% | 0.0% |  |
| *serpentine* | **TRiP** | **65104** | **<u>15.4%</u>** | **18.2%** | **Bent or broken tarsals** |
| *vermiform* | TRiP | 57188 | <u>0.0%</u> | N/A | Died as prepupae |
| **Miscellaneous** |  |  |  |  |  |
| *Amalgam* | TRiP | 33416 | <u>14.8%</u> | 0.0% |  |
|  | VDRC | 22944 | 45.2% | 0.0% |  |
| *CG5758* | TRiP | 57808 | 33.3% | 0.0% |  |
| *slowdown* | VDRC | 106464 | 45.4% | 3.6% |  |
| *CG10960* | TRiP | 34598 | 45.8% | 0.0% |  |
| *CG12026* | VDRC | 42480 | <u>42.7%</u> | 0.6% |  |
| *grainy head* | **TRiP** | **42611** | **<u>37.8%</u>** | **5.6%** | **Short, fat tarsals** |
| *Ecdysone-inducible gene E3* | VDRC | 16402 | 40.0% | 0.0% |  |

Lines listed include both TRiP and VDRC (long hairpin) lines, with stock numbers shown. Lines exhibiting a significant number of malformed legs are indicated in bold. Percentages of adult flies carrying the Gal4 driver (determined by absence of curly wings) is indicated; values found by chi-square test to be significantly lower than 50% are underlined. All RNAi lines were crossed with virgin females carrying the *Dll*-Gal4 driver and a UAS-*Dicer* construct.

In control experiments, we drove the expression of *br*-RNAi in distal leg segments and observed no adults; +4-hour leg discs from *Dll>br-RNAi* animals revealed defects in leg elongation similar to that observed in *br^5^* mutant leg discs (Fig. S2 B, C). Distal leg expression of *Sb-RNAi* resulted in 18% of the animals displaying a malformed leg phenotype characterized by shorter, fatter distal tibia and tarsal segments (Fig. 6B). This malformed phenotype is similar to that observed in a hypomorphic *Sb* allele (Fig. 6C). These experiments validate the use of *Dll-GAL4* to reduce gene expression of *br*-induced genes during late larval and prepupal leg morphogenesis.

**Fig. 6.**
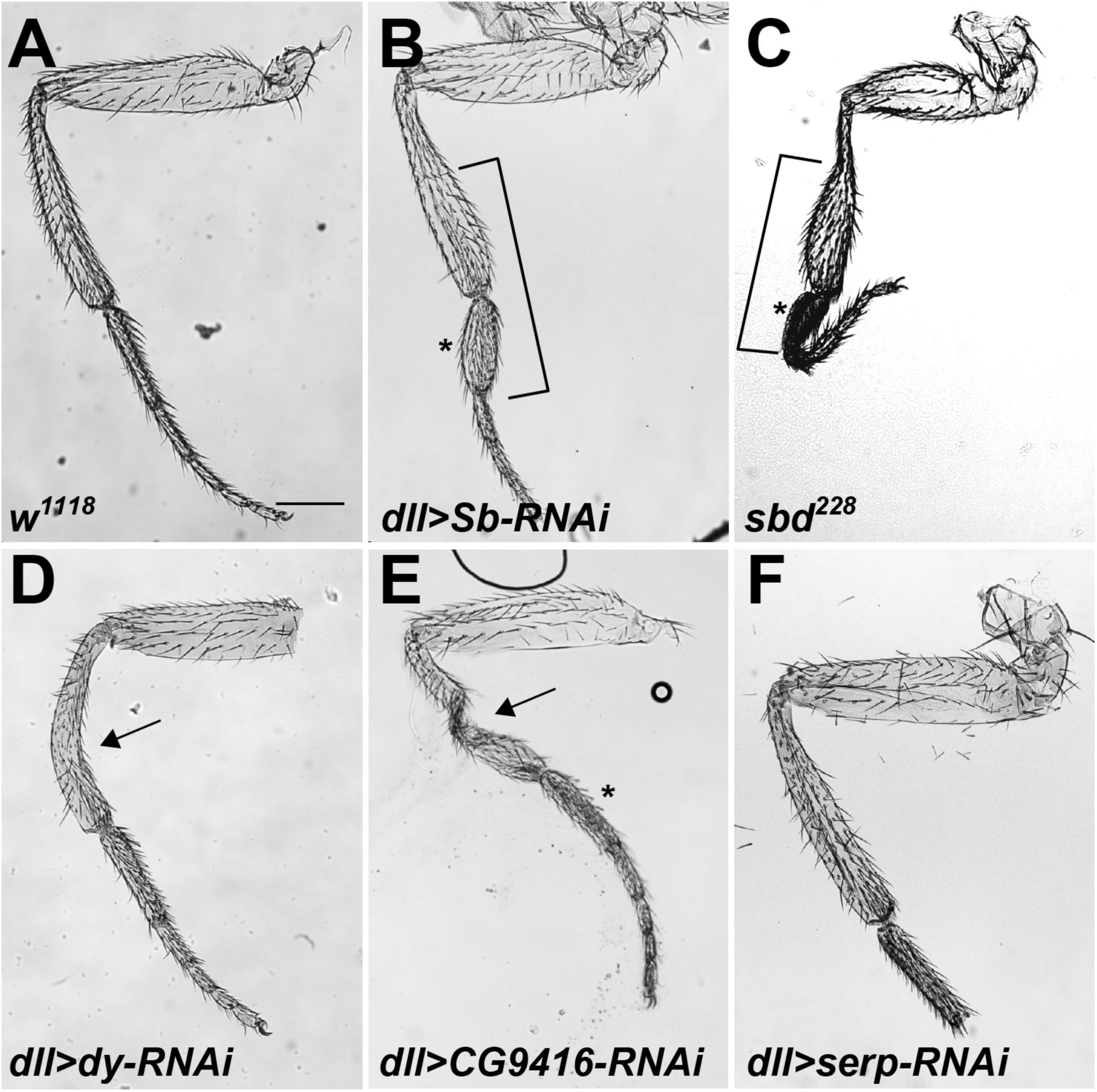
Driving RNAi against some *br*-induced genes in distal leg segments leads to malformed adult legs. Brightfield photomicrographs of adult legs from *w^1118^* (A), *Dll>Sb-RNAi* (B), *Sb^Ebr228^* (C), *Dll>dy-RNAi* (D), *Dll>CG9416-RNAi* (E), and *Dll>serp-RNAi* (F). Note that the *Dll>Sb-RNAi* (B) has a shorter and fatter distal tibia and tarsal segment similar to that observed in the homozygous *sbd* mutant (brackets) (C). Malformed phenotypes include bends or kinks in the tibia (arrows), and misshapen tarsal segments (asterisks). *Dll>serp-RNAi* flies (F) show a high penetrance of missing distal tarsal segments. Scale bar equals 200 um.

We next drove RNAi against 27 *br*-induced genes using both long-hairpin and short-hairpin *UAS-RNAi* lines with the *Dll*-*GAL4* driver. A total of 32 RNAi lines were tested, and phenotypes similar to those reported in screens by Ward *et al*., including bent tibias and poorly proportioned tarsals (Ward *et al*., 2003), were observed in several of the lines (Table 2 and Fig. 6). In total, five lines exhibited malformed legs at a rate greater than 4%, including two lines from the protease and protease inhibitor group, *CG9416* and *Sb*, and two lines knocking down chitin-related genes, *dusky* (*dy*) and *serp*. The final line carried a construct against *grainy head* (*grh*). In all cases, straight winged flies that carried the *GAL4* driver showed significantly more leg defects than curly-winged flies from the same cross that did not carry the GAL4 driver (*p* < 0.001). The phenotypes observed in each of these crosses differed (Fig. 6): in those from the *dy* line, flies exhibited slightly curved tibias, while flies from the *CG9416* line exhibited more severely kinked tibias. *serp* line flies frequently showed bent or missing tarsal segments, while the tarsal segments in *Sb* and *grh* flies were often misshapen or misproportioned (Fig. 6B and data not shown). Variation in phenotype penetrance and expressivity also existed between crosses with lines carrying constructs against the same genes (Table 2). For example, bent tibias were observed in 38% of flies expressing a short-hairpin construct against *CG9416*, while flies expressing a long-hairpin construct did not display this phenotype.

Several lines produced fewer of the straight-winged RNAi-expressing flies than expected, suggesting some lethal effect. In total, eleven crosses targeting *br*-induced genes resulted in significantly less than 50% of straight wing flies (Table 2). These include four crosses (*CG9416*, *grh*, *Sb*, and *serp*) that exhibited malformed legs at a rate greater than 4%, as well as two crosses (*vermiform* [*verm*] and *Enhancer of split m*α [*E(spl)m*α]) that produced no RNAi-expressing straight wing flies that could be scored for the malformed leg phenotype. Individuals expressing RNAi against *E(spl)m*α failed to undergo larval molts, appearing to remain as first instar larvae for several days before dying (data not shown). In contrast, individuals expressing RNAi against *verm* pupariated, with the majority of the animals pupating, but without everted legs. After about a day following pupation, there was noticeable necrotic tissue in the areas where the legs, the antennae, and wing margins form in wild type pupae (Fig. 7B). When we dissected +4 hr leg discs from *Dll>verm-RNAi* prepupae, we found that they exhibited an elongation defect that was more severe than what was observed in *br^5^* mutants or *br* RNAi knockdown flies (compare Fig. 7D to Fig. S2 B and C). To investigate whether the majority of the *br^5^* mutant phenotype is due to loss of *verm*, we drove UAS-*verm* in *br* RNAi imaginal discs (using *Dll-GAL4*). Neither expression of *verm* (nor its fellow chitin deacetylase, *serp*), was able to rescue *br* RNAi flies to adulthood (7 independent experiments with *UAS-verm* with n = 409 adults eclosing and 4 independent experiments with *UAS-serp* with n = 291 adults eclosing).

**Fig. 7.**
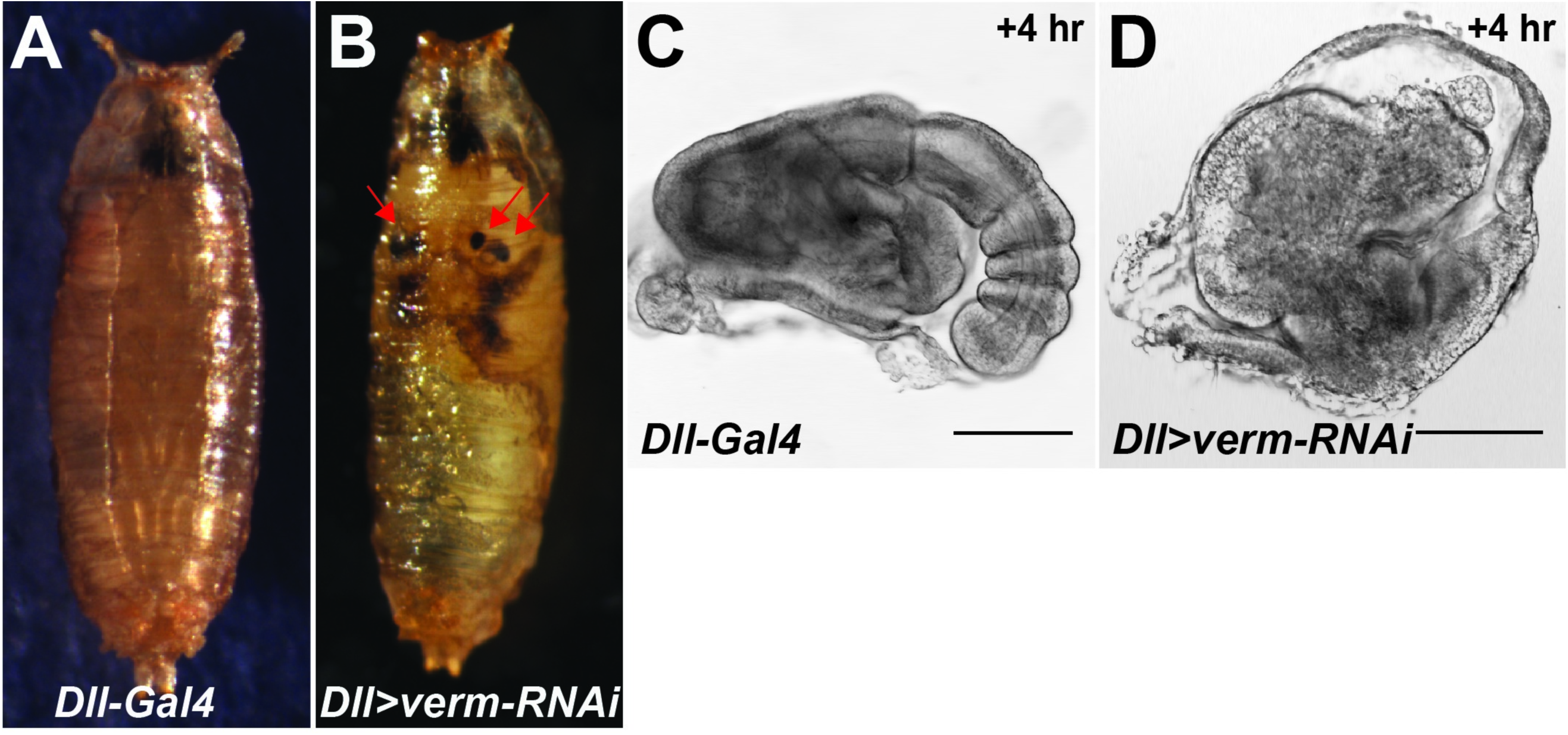
Distal leg expression of *verm-RNAi* leads to severe defects in leg morphogenesis. (A, B) Color photomicrographs of *Dll-Gal4* (A) and *Dll>verm-RNAi* (B) mid-stage pupae. Note the lack of elongated legs in the *Dll>verm-RNAi* animal that instead shows necrotic tissue in the areas where the legs should evert from (arrows). (C, D) Brightfield photomicrographs of leg imaginal disc from *Dll-Gal4* (C) and *Dll>verm-RNAi* (D) +4 hr prepupae showing little distal elongation and almost no segmentation in the *verm-RNAi* disc. Scale bar equals 100 um.

## Discussion

In this study, we have demonstrated that Br is required for many of the tissue-level events required for leg morphogenesis during the first several hours of metamorphosis in *Drosophila*, and have identified a collection of genes regulated by *br* that may help explain this developmental process. Many previous studies have demonstrated that leg morphogenesis requires ecdysone signaling (Mandaron, 1971; Fristrom *et al*., 1973), cell shape changes and rearrangements (Condic *et al*., 1991), and proteolysis of the ECM (Diaz-de-la-Loza *et al*., 2018). Here we show that leg discs in *br^5^* mutant animals fail to fully degrade their ECM (Fig. 2), and that even if the ECM is degraded by exogenous trypsin they fail to fully elongate (Fig. 3), suggesting that the underlying cell shape changes and rearrangements are also defective. Anisometric cell shapes and wider tarsal segments in +4 hour leg discs from *br^5^* mutant animals support this suggestion (Fig. 3). Consistent with all of these observations, RNA-Seq comparisons between *br^5^* and *w^1118^* leg imaginal discs at the onset of morphogenesis indicate that *br* regulates the expression of several potential proteases and chitin modifying genes. RNAi-based functional analyses confirm a requirement for some of these *br*-induced genes in leg imaginal disc morphogenesis (Figs. 6 and 7). Furthermore, bioinformatic analyses of *br*-regulated genes suggests that *br* is required to maintain robust metabolism in imaginal discs when glycolysis is otherwise downregulated in the larval tissues.

### Bioinformatic analyses of gene functions suggest a role for Br in maintaining metabolic activity during metamorphosis

The amorphic *br^5^* allele leads to failure of imaginal disc development, and, as expected, several development-related biological process GO terms were enriched in *br*-induced genes, including “developmental process,” “anatomical structure development,” and “organ morphogenesis.” Somewhat surprisingly, the enriched GO terms also included glycolysis-related metabolic terms, including “pyruvate metabolic process,” “ATP generation from ADP,” and “glycolytic process.” In support of these findings, KEGG pathway analysis identified Glycolysis/Gluconeogenesis as the most significantly enriched pathway from these genes. More interestingly, neither analysis revealed enrichment of glycolysis-related metabolic genes in the comparison of *w^1118^* leg discs from −18hr to 0hr (neither up- or down-regulated at 0 hr). This finding contrasts with previous microarray studies in both *Drosophila melanogaster* and the silkworm *Bombyx mori* showing downregulation of glycolytic and other metabolic genes in whole animals at pupariation (White *et al*., 1999; Tian *et al*., 2010). Taken together, these results suggest that while larval tissues are shutting down metabolism (Merkey, 2011), the imaginal discs retain metabolic activity throughout the transition to prepupa, and that this metabolic program requires Br. This energy requirement could be needed for the cell shape changes and rearrangements that drive leg morphogenesis. We were therefore surprised to find ATP levels were higher in *br^5^* mutant discs than in *w^1118^* discs at 4 hr after pupariation (Fig. S1), but this result may instead reflect the reduced energy consumption of *br^5^* mutant leg discs that have already arrested morphogenesis by 4 hr APF.

### *br*-dependent extracellular matrix remodeling plays a critical role in leg development

Our results suggest that *br* plays a central role in remodeling the ECM in leg imaginal discs during metamorphosis. We show that Collagen IV is not cleared from the basal ECM in *br^5^* mutant prepupal leg discs (see Fig. 2). In addition, four of the six genes that resulted in malformed adult legs when downregulated by RNAi (*dusky, serpentine*, *Stubble,* and *vermiform*) have been proposed to interact with the apical ECM, while another *(grainy head*) is known to regulate ECM genes (Pare *et al*., 2012; Yao *et al*., 2017). *Stubble* encodes a serine protease whose endopeptidase domain is required for normal leg development (Appel *et al*., 1993; Hammonds and Fristrom, 2006). Stubble protein has been shown to be required for degradation of the Dumpy protein, an apical ECM component that links the epithelial cells to the cuticle (Maria-del-Carmen *et al*., 2018). In the absence of this degradation, imaginal discs fail to elongate. *serpentine* and *vermiform* were identified as genes encoding chitin deacetylases involved in organization of the larval tracheal apical ECM (Luschnig *et al*., 2006). These genes are also found in the larval cuticle, where they are apically secreted and necessary for the formation of a stable matrix (Pesch *et al*., 2015; Pesch *et al*., 2016). The *dusky* gene encodes a zona pellucida protein that is expressed in epithelial tissues across *Drosophila* development, most notably in the developing wing disc during wing elongation (Roch *et al*., 2003; Jazwinska and Affolter, 2004; Ren *et al*., 2005). Zona pellucida proteins are membrane-anchored proteins that interact with the ECM; a study in embryonic denticles showed localization to specific apical subcellular domains, where the proteins organize and modify the ECM to control cell shape changes (Fernandes *et al*., 2010). *grainy head* encodes a transcription factor that regulates genes involved in cuticle formation (Bray *et al*., 1989; Dynlacht *et al*., 1989; Bray and Kafatos, 1991). *grainy head* mutants display defective ECM phenotypes in the head skeleton (Nusslein-Volhard *et al*., 1984; Bray and Kafatos, 1991), trachea (Hemphala *et al*., 2003) and in wound repair (Mace *et al*., 2005). ChIP data suggests that Grainy head also regulates some of the other leg morphogenesis genes identified in this study, including *CG9416*, *serpentine, vermiform,* and *Stubble* (Pare *et al*., 2012; Yao *et al*., 2017).

The connection that most of these genes have with the ECM is consistent with the known role of the ECM in imaginal disc elongation; the ECM provides a constraining force to imaginal discs (Pastor-Pareja and Xu, 2011), and both the apical and basal ECMs must be degraded before elongation (Maria-del-Carmen *et al*., 2018). Our results suggest that *br* provides a key regulatory mechanism for these processes, and that they underlie a large part of the *br^5^* phenotype. *br* does not appear to be wholly responsible for the proteolysis of the ECM at metamorphosis, however. Notably absent from our list of *br*-regulated genes are the matrix metalloprotease genes *Mmp1* and *Mmp2*, which are required for breakdown of the basal extracellular matrix during disc eversion (Srivastava *et al*., 2007), although both of these genes are developmentally upregulated at the onset of metamorphosis in leg discs (Table S4).

### *broad* is dynamically regulated before and during metamorphosis

The *Broad-Complex* was initially described as an early-response gene in the ecdysone-induced transcriptional cascade (Chao and Guild, 1986; DiBello *et al*., 1991), but subsequent experiments suggested that the regulation of *br* by ecdysone both in whole animals and in imaginal discs is more complex and involves an isoform switch from *Z2/Z3* to *Z1* at pupariation. Specifically, studies in the salivary glands and whole animals showed a switch to *Z1* upon pupariation (Andres *et al*., 1993; Huet *et al*., 1993), and imaginal discs cultured in the presence of ecdysone resulted in persistence of the *Z1* transcript even after the other *br* isoforms were no longer detected (Bayer *et al*., 1996). Our northern blot analysis confirms this isoform switch at the onset of metamorphosis in imaginal discs, and also reveals that *br Z2/Z3* is expressed earlier in the discs than would be predicted by whole animal blots (Fig. 5A). This latter observation is somewhat paradoxical considering that the *br*-induced genes are a reasonably good subset of the developmentally-induced genes revealed by RNA seq comparisons of leg discs from −18 hr to 0 hr (Fig. 4). This raises the possibility that many of these *br*-induced genes are also regulated in a temporal fashion, perhaps by ecdysone/EcR/Usp directly. Future experiments should be aimed at exploring the more complex regulation of these genes, particularly those that we have identified as playing important roles in driving leg morphogenesis. It is also interesting that Br appears to be responsible for controlling early prepupal developmental events (proteolysis of the ECM, cell shape changes and rearrangements) when the Z2 isoform is ostensibly “off.” We believe that this is driven by perdurance of the Br protein. In support of this, Emery *et al*. (Emery *et al*., 1994), demonstrated that levels of the Br-Z2 protein remained strong in the imaginal discs until between 4 hr and 8 hr after pupariation, despite the decrease in transcription by 0 hr. Finally, it is worth noting that we identified a large number of *br*-repressed genes in our RNA seq experiment, raising the possibility that derepression of *br*-repressed genes may also contribute to normal leg morphogenesis. Our functional studies in which we extended *br-Z2* expression through a heat inducible promoter resulted in complete lethality, with half of the animals dying prior to pupation. The terminal leg phenotypes in the prepupal lethal animals revealed that development was normal until 4-7 hours after pupariation. For those animals that pupated, their legs and wings were short, the heads were often malformed and they died soon after pupation. Taken together, these findings are consistent with a model in which perdurance of Br is sufficient to positively regulate genes needed for early prepupal leg development, at which point reduced Br results in derepression of a second set of genes that positively affects later prepupal and pupal leg development.

## Materials and Methods

### Fly stocks

All *Drosophila* stocks were maintained on media consisting of corn meal, sugar, yeast, and agar in incubators maintained at a constant temperature of 25° or at room temperature. *w^1118^*, *y br^5^*, *distalless* (*dll*)*-Gal4*, *apterous (ap)-Gal4*, *P{UAS-Dcr-2.D}1 w^1118^*, and the Transgenic RNAi Project (TRiP) lines (“short-hairpin” RNAi lines; https://fgr.hms.harvard.edu) were obtained from the Bloomington *Drosophila* Stock Center (Bloomington, IN). “Long-hairpin” RNAi lines were obtained from the Vienna *Drosophila* RNAi Center (VDRC, Vienna, Austria; (Dietzl *et al*., 2007)). *UAS-serp* was obtained from Stefan Luschnig (University of Muster). *UAS-Verm/CyO* was obtained from Christos Samakovlis (Stockholm University). *vkg-GFP* (Flytrap; (Buszczak *et al*., 2007)) was obtained from Sally Horne-Badovinac (University of Chicago). *w; hs-Z2 (CD5-4C); hs-Z2 (CD5-1)* was obtained from Cindy Bayer (University of Central Florida). *Loxl-1-RNAi* (stock 11335R) was obtained from the National Institutes of Genetics Fly stock collection (Kyoto, Japan). *dll-Gal4* was balanced with *CyO, P{w^+^, Dfd-EYFP}* (Le *et al*., 2006). *w^1118^* was used as the wild type control, unless otherwise noted.

### Fly staging, dissection and photography of live leg imaginal discs

*w^1118^* and *y br^5^*/*binsn* flies were staged on food supplemented with 0.05% bromophenol blue as described in (Andres and Thummel, 1994). *w^1118^*and *y br^5^*/Y mutant animals were selected (mutant males were selected using the *yellow* marker) at −18 hr (blue gut larvae), −4 hr (white gut larvae), 0 hr (white prepupae), +2 hr and +4 hr relative to puparium formation. Leg imaginal discs were dissected in Phosphate Buffered Saline (PBS), and then transferred to fresh PBS. Brightfield photomicrographs were captured within 5 minutes on a Nikon Eclipse 80i microscope equipped with a Photometrics CoolSNAP ES high performance digital CCD camera using a Plan APO 10X (0.45NA) objective. For the examination of collagen integrity in prepupal leg discs, we crossed y*br^5^/Binsn* virgins to *w^1118^, vkg-GFP* males, and crossed F_1_y*br^5^/w^1118^; vkg-GFP/+* females to *w^1118^* males. We selected *ybr^5^/Y; vkg-GFP/+* and *w^1118^/Y; vkg-GFP/+* white prepupae based upon the *y* phenotype and aged them at 25°. *ybr^5^* prepupae were confirmed to contain the *br^5^* allele based upon pupal morphology. We dissected leg imaginal discs at the indicate time points in PBS, mounted them live in mounting media (90% glycerol, 100 mM Tris pH 8.0, 0.5% n-propyl-gallate) and imaged them sequentially with brightfield microscopy and wide-field fluorescence microscopy on a Nikon Eclipse 80i microscope equipped with a Photometrics CoolSNAP ES high performance digital CCD camera using a Plan APO 20X (0.75NA) objective. All images were captured with identical settings (light or fluorescence intensities, and exposure times). On average 3-4 discs were imaged from each prepupa, and at least 7 different prepupae were imaged at each time point. All digital images were cropped and adjusted for brightness and contrast in Adobe Photoshop (version CC 2018, San Jose, CA) or ImageJ (version 1.51r, National Institutes of Health, Bethesda, MD), and Figures were compiled using Adobe Illustrator (version CC 2018).

### Trypsin experiments

*w^1118^* and *br^5^* white prepupae were dissected in PBS to isolate three leg imaginal discs from the same animal. These imaginal discs incubated on 3-well depression slides in either PBS, PBS plus 0.025% trypsin (Gibco, 25200-056) or PBS plus 0.0025% trypsin for 15 minutes at room temperature, at which point they were immediately imaged by brightfield microscopy on a Nikon Eclipse 80i microscope equipped with a Photometrics CoolSNAP ES high performance digital CCD camera using a Plan APO 10X (0.45NA) objective. In total, 21 *w^1118^*and 17 *br^5^* animals were dissected and imaged.

### Immunostaining

Imaginal discs were hand dissected from *w^1118^*or *br^5^* +4 hr prepupae in fresh PBS, and fixed immediately in 4% paraformaldehyde for 20 minutes. The following antibodies were used at the given dilutions: rat anti-DEcadherin (clone DCAD2 from Developmental Studies Hybridoma Bank at the University of Iowa) 1:25 and Donkey anti-rat Cy2 (Jackson ImmunoResearch Laboratories, West Grove, PA) at 1:400. Confocal images were acquired on a Leica SPE laser scanning confocal microscope using an ACS APO 40X (1.15 NA) oil immersion lens. All digital images were cropped and adjusted for brightness and contrast in ImageJ (version 1.51s, National Institutes of Health, Bethesda, MD). Figures were compiled using Adobe Illustrator (version CC 2018).

### RNAseq and data analysis

In order to control for potential genetic effects from the autosomes and Y chromosome, we crossed *ybr^5^/Binsn* females to *w^1118^* males and then crossed the resulting *ybr^5^/w^1118^*females with *w^1118^* males. This cross produced *ybr^5^/Y* and *w^1118^/Y* males that at a population level had identical autosomes and Y chromosomes. Since we selected the *br^5^* animals based upon the *yellow* cuticular phenotypes, we tested to make sure that *y* and *br* did not recombine apart to any significant degree. *y* and *br* are reported to map 0.2 cM apart on the X chromosome (Gatti and Baker, 1989), and in two separate experiments we did not detect any recombination between these genes (*n* > 200). After RNA sequencing, we identified the *br^5^* mutation as a C to T transition at position 1654571 in Genbank AE014298 (X chromosome of *Drosophila melanogaster*), converting a conserved histidine in the zinc finger of the Z2 isoform into a tyrosine. We found that all the reads through this interval had the mutation in the *br^5^* samples, whereas none of the reads from *w^1118^*samples had the mutation.

Third instar *ybr^5^/Y* and *w^1118^/Y* larvae were staged on food supplemented with 0.05% bromophenol blue. Blue gut larvae (−18 hr) and white prepupae (0 hr) were selected and leg imaginal discs were hand-dissected in PBS. Triplicate independent samples were obtained for *ybr^5^/Y* and *w^1118^/Y* white prepupae and for *w^1118^/Y* −18 hour larvae. Total RNA was isolated using TriPure (Roche, Indianapolis, IN) and then purified over RNAeasy columns (Qiagen, Valencia, CA). ∼5 μg of total purified RNA was obtained for each sample, and ∼1 ug of each sample was provided to the Genome Sequencing Core at the University of Kansas for library preparation using the TruSeq stranded mRNA kit (Illumina, San Diego, CA). Single Read 100 (SR100) was performed on a single lane of an Illumina HiSeq 2500 (Genome Sequencing Core, KU).

The quality of the raw RNAseq reads was visually confirmed using FastQC (version 0.11.5, (Andrews, 2010)), and both low quality data and adaptor sequences were removed using Trimmomatic (version 0.36, (Bolger *et al*., 2014)). The number of remaining reads per sample averaged 18.1 million (range = 12.6-22.8), and all reads were at least 50-nt in length. Filtered reads were mapped to the *D. melanogaster* reference genome (Release 5.3) using TopHat (version 2.1.1, (Trapnell *et al*., 2009; Kim *et al*., 2013)). Default parameters were employed, with the addition of the “--no-novel-juncs” flag to use the gene annotation as provided, and the “--library-type fr-firststrand” flag since the RNAseq data derives from the Illumina TruSeq stranded mRNA kit. On average 86.2% of the reads mapped to the reference across samples (range = 83.1-88.5). Genotypes were compared, and differentially-expressed genes were identified using the Cufflinks pipeline (version 2.2.1, (Trapnell *et al*., 2010; Roberts *et al*., 2011; Trapnell *et al*., 2013)). Specifically, we employed the “cuffquant” and “cuffdiff” routines using default paramaters, adding the “-b” flag to run a bias correction algorithm that improves expression estimates (Roberts *et al*., 2011).

### Bioinformatic analyses

Gene ontology analysis was performed using the Gene Ontology Consortium’s (http://www.geneontology.org) Gene Ontology Enrichment tool (The Gene Ontology Consortium. 2015). Significantly differentially expressed genes showing ≥ 1.5-fold-change between *w^1118^* 0 hr and *br^5^* 0 hr samples and at least 5 FPKM in the sample showing higher expression (*w^1118^* for *br*-induced genes, *br^5^ br*-repressed genes) were used as input into the tool. In a second experiment, significantly differentially expressed genes showing ≥ 1.5-fold-change between *w^1118^* −18 hr and 0 hr samples and at least 5 FPKM in the sample showing higher expression (*w^1118^* 0 hr for developmentally-induced genes and *w^1118^* −18 hr for developmentally*-*repressed genes) were used. The same data sets were used for the KEGG pathway analysis using WebGestalt’s (http://www.webgestalt.org) Over-Representation Analysis (ORA) (Wang *et al*., 2017).

### ATP measurement

Leg and wing imaginal discs were dissected from +4 hr *w^1118^* and *br^5^* mutant prepupae in PBS and immediately homogenized in 100 μl of lysis buffer (6M guanidine HCl; 100 mM Tris pH 7.8, 4 mM EDTA) on ice and frozen. After all the samples were collected, they were thawed to 4°C, and a10 μl aliquot was taken for protein measurement using a Bradford Assay (Bio-Rad, Hercules, CA). The remaining sample was boiled for 5 min, spun at 13,000 rpm in a refrigerated microcentrifuge for 3 min and then 10μl of the supernatant was diluted into 90 μl of dilution buffer (25mM Tris pH7.8, 100 μM EDTA). This sample was further diluted 10-fold in dilution buffer and 10 μl of this sample was used for ATP quantification using the ATP Determination Kit (Molecular Probes A22066, Eugene, OR) according to the manufacturer’s protocol using a BioTek Synergy HT plate reader. Each biological sample contained approximately 13 wing discs and 35 leg discs. Triplicate samples were processed and three technical replicates of each biological sample was assayed, along with a dilution series of ATP from 1 nm to 1 μM. Statistical analysis was performed using a type 3 ANOVA with protein level and treatment as factors.

### Northern blot analysis

Progeny from a cross of *y br^5^/Binsn* X *Binsn/Y* were staged on standard *Drosophila* media supplemented with 0.05% bromophenol blue as described in (Andres and Thummel, 1994). Total RNA was isolated by direct phenol extraction from leg imaginal discs dissected from staged *y br^5^/Y* and *Binsn/Y* males. Approximately 5 ug of total RNA per sample were separated by formaldehyde agarose gel electrophoresis and transferred to a nylon membrane. The membrane was hybridized and stripped as described in (Karim and Thummel, 1991). Generation of probe fragments for *br* (*BR-C* core) and *rp49* is described in (Andres and Thummel, 1994). Specific probes were labeled by random priming of gel-purified fragments (Stratagene).

### Functional analyses

To amplify potential RNA interference in long-hairpin lines, we crossed a *UAS-Dicer* transgene onto the *Gal4* lines used to drive the expression of the gene-specific double stranded RNA to produce *P{UAS-Dcr-2.D}1, w^1118^; Dll-GAL4 /CyO, dfd-YFP* and *P{UAS-Dcr-2.D}1, w^1118^; ap-GAL4 /CyO, dfd-YFP.* For consistency, we also used these recombined lines when crossing to short-hairpin RNAi lines. Virgin females of these genotypes were then mated to *UAS-RNAi* males. The vials were kept in incubators maintained at a constant temperature of 25°C. The adults were transferred twice to new vials and newly eclosing F_1_ flies were separated by phenotype and examined for malformed third legs each day for a total of 8 days per vial. We considered an animal to be malformed if it displayed malformation in at least one leg, and defined a leg as malformed if any femur, tibia or tarsal segment was bent, twisted, missing, or was excessively short and fat.

To control for background effects that may contribute to the appearance of malformed legs, we crossed *UAS-Dcr1, w^1118^; Dll-GAL4 /CyO, dfd-YFP* virgin females with males from the *w^1118^*line and a *loxl1-*RNAi line and examined the offspring for malformed legs. Curly-winged offspring from RNAi crosses, which carry the *UAS-RNAi* hairpin construct, but not the GAL4 driver, were also examined. Flies carrying malformed legs did not exceed 1% of all flies in either control cross, and exceeded 1% among curly-winged offspring from only 2 RNAi crosses. In neither of these two crosses (carrying RNAi hairpin constructs against *CG7447* and *E[spl]m4*) did flies carrying malformed legs exceed 2% of all curly-winged flies, and neither of these crosses were among those in which a significant number of straight-winged flies carried malformed legs. For crosses in which we examined pre-adult stages expressing RNAi, individuals carrying the *Dll*-*GAL4* driver were identified by the lack of *dfd-YFP* expression.

### *br-Z2* overexpression studies

Approximately 50 *HS-br-Z2* or *w^1118^*late larvae were placed into an empty fly vial with a piece of moist whatman paper in the bottom of the vial. The vial was heat shocked in a 37° incubator for 60 minutes, after which any animals that had pupariated were removed. The vial was moved to a 25° incubator for 6 additional hours at which point all prepupae were removed to a food vial for further development (also at 25°). 18 hours later the prepupae were scored to determine if they had pupated. The animals were then left at 25° and scored for eclosing 4 days later. In some experiments, *HS-br-Z2* or *w^1118^* were collected as they pupariated, aged 4 hours at 25° and dissected to examine their leg imaginal discs. Another subset was dissected at approximately 16 hours after pupariating to examine the terminal leg imaginal disc phenotypes. All dissections, microscopy and image preparation were conducted as described above.

### Adult specimen preparations

Adult legs were dissected from the third thoracic segment in PBS, cleared in 10% KOH overnight, and mounted in Euparal (Bioquip, Gardena, CA) on microscope slides. Images of adult leg cuticles were captured on a Photometrics CoolSNAP ES high performance digital CCD camera with a Nikon Eclipse 80i microscope. All digital images were cropped and adjusted for brightness and contrast in ImageJ (version 1.51r, National Institutes of Health, Bethesda, MD) and Figures were compiled using Adobe Illustrator (version CC 2018).

## Acknowledgements

We thank Stefan Luschnig, Christos Samakovlis, Cindy Bayer, Sally Horne-Badovinac, the Bloomington *Drosophila* Stock Center, and the Vienna *Drosophila* RNAi Center for fly stocks. We thank Taybor Parker, David Davido, and Rob Unckless for help with the ATP measurement and analysis. The DCAD2 antibody developed by T. Uemura was obtained from the Developmental Studies Hybridoma Bank, created by the NICHD of the NIH and maintained at The University of Iowa, Department of Biology, Iowa City, IA 52242. We also thank Rebecca Spokony, Laurie von Kalm, Jason Tennessen, and members of the Ward lab for helpful discussion about the project and manuscript.

## Funding

This work was supported by the College of Liberal Arts and Sciences at the University of Kansas (CR). RNA sequencing was supported by a KU genome sequencing core users grant (RW). The sequencing core is supported by National Institutes of Health grant P20 GM103638. Analysis of RNA sequencing data (by SM) was aided by infrastructure purchased via the Kansas IDeA Network of Biomedical Research Excellence (National Institutes of Health grant P20 GM103418). The funders had no role in study design, data collection and analysis, decision to publish, or preparation of the manuscript.

## Competing Interests

The authors have declared that no competing interests exist.

## Data Availability

All RNA seq datasets are available from the Gene Expression Omnibus. The project accession number is GSE140248, and the project title is “Genome-wide differences in RNA expression in Drosophila melanogaster leg imaginal discs based on time and presence/absence of broad-based gene regulation.”

## Figure Legends

**Fig. S1. Comparison of luciferase ATP assay luminescence normalized to protein in *br^5^* and *w^1118^* leg and wing imaginal discs.** Leg and wing imaginal discs were pooled into three biological replicates for both wild type and mutant lines and ATP levels were assessed using a luciferase-based assay and normalized to protein levels for each sample. Lines represent average values from three technical replicates (shown as dots) from each sample. We performed a type 3 ANOVA with protein level and treatment as factors. ATP levels were significantly different both when comparing across treatment (*w^1118^* vs. *br^5^*; P=0.03) and when comparing between samples based on protein level (P=0.0000496).

**Fig S2. Distal leg expression of *br-RNAi* recapitulates *br^5^* leg phenotypes.** Brightfield photomicrographs of leg imaginal discs from +4 hr *Dll-Gal4* (A), *Dll>br-RNAi* (B) and *br^5^* (C) prepupae. Discs from *Dll>br-RNAi* animals show a similar lack of elongation and segmentation defects as seen in the *br^5^* mutant discs. Scale bar equals 100 um.

